# Modelling brain representations of abstract concepts

**DOI:** 10.1101/2021.06.02.446744

**Authors:** Daniel Kaiser, Arthur M. Jacobs, Radoslaw M. Cichy

**Affiliations:** Mathematical Institute, Department of Mathematics and Computer Science, Physics, Geography, Justus-Liebig-Universität Gießen; Center for Mind, Brain and Behavior (CMBB), Philipps-Universität Marburg and Justus-Liebig-Universität Gießen; Department of Psychology, University of York; Department of Education and Psychology, Freie Universität Berlin; Center for Cognitive Neuroscience Berlin, Freie Universität Berlin; Berlin School of Mind and Brain, Humboldt-Universität zu Berlin; Bernstein Center for Computational Neuroscience Berlin

## Abstract

Abstract conceptual representations are critical for human cognition. Despite their importance, key properties of these representations remain poorly understood. Here, we used computational models of distributional semantics to predict multivariate fMRI activity patterns during the activation and contextualization of abstract concepts. We devised a task in which participants had to embed abstract nouns into a story that they developed around a given background context. We found that representations in inferior parietal cortex were predicted by concept similarities emerging in models of distributional semantics. By constructing different model families, we reveal the models’ learning trajectories and delineate how abstract and concrete training materials contribute to the formation of brain-like representations. These results inform theories about the format and emergence of abstract conceptual representations in the human brain.

## Introduction

The use of conceptual knowledge is one of the foundations of human intelligence. On the neural level, concepts are represented in a complex network of brain regions (Binder et al., 2009). Fueled by novel computational models of distributional semantics, researchers have recently started to unravel the format of concept representations in this neural network. By harnessing linguistic co-occurrence statistics, these models not only capture representations of concepts from written and spoken language (Deniz et al., 2019; Huth et al., 2016; Just et al., 2010; Mitchell et al., 2008), but also predict representations of novel concepts (Pereira et al., 2018).

However, these recent advances in understanding the representations of conceptual knowledge largely hinge on the study of concrete concepts. And although some of the previous studies (e.g., Huth et al., 2016, Pereira et al., 2018) have included both concrete and abstract words, they probed representations across the two, and therefore, could not reveal how specifically abstract concepts are represented. Only few studies have explicitly investigated how abstract concept representations are organized (Anderson et al., 2014, 2017; Vargas & Just, 2020; Wang et al., 2018a). To date, key questions about the emergence and the format of these representations remain heavily debated (Borghi et al., 2017).

Here, we model brain representations that support the activation and contextualization of abstract concepts. We recorded fMRI while participants were tasked with embedding abstract nouns into a background context. By relating brain activations during this task to targeted models of distributional semantics, we shine a new light on the format of abstract concept representations in the human brain.

## Results and Discussion

In an fMRI experiment, we visually presented 61 abstract German nouns (see Materials and Methods). Participants (n=19) read these words and silently embedded them into a coherent story that they were developing around a prespecified contextual background (Figure 1a). This task was chosen to be engaging while ensuring a sufficiently deep level of processing. Here, participants were required to retrieve the meaning of the words and use those meanings, integrating them with a complex ongoing stream of thought.

**Figure 1.**
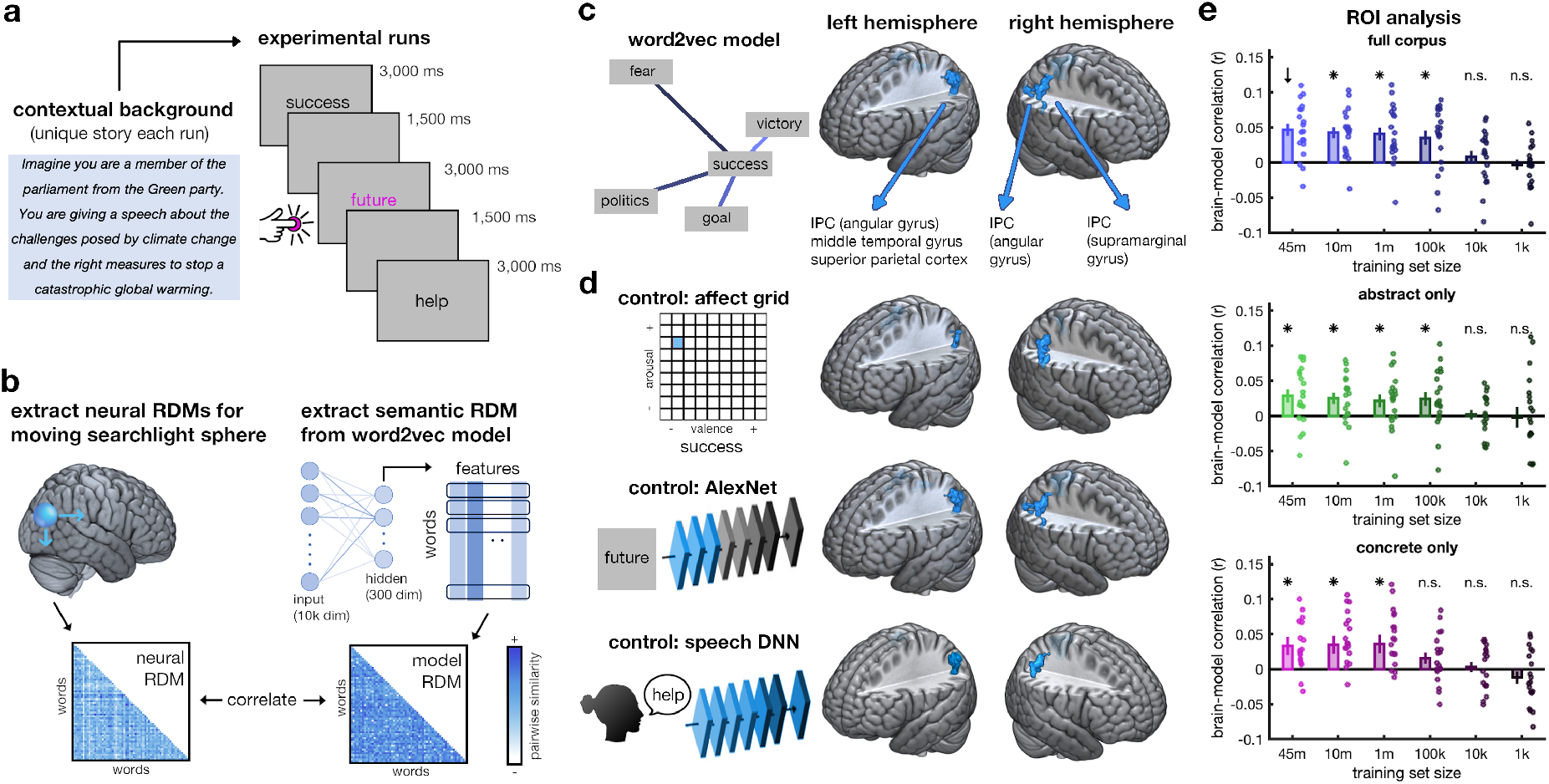
Representation of abstract concepts in parietal cortex. **a)** Participants completed 10 runs of fMRI recordings. Before each run, they established a unique background context in their mind and were then asked to silently narrate a story of their own making that included the subsequently presented 61 abstract words, in random order (all German word stimuli in Supplementary Information, Table S1). **b)** In a searchlight analysis, representational dissimilarity matrices (RDMs) were extracted (i) from the brain data, by pairwise correlations among localized activity patterns, and (ii) from a word2vec model of distributional semantics, by pairwise correlations among hidden-layer activations. **c)** Correlating the neural and model RDMs revealed clusters in bilateral inferior parietal cortex (IPC), primarily covering the angular gyrus. Brain maps are thresholded at p_voxel_<0.001 (uncorrected) and p_cluster_<0.05 (FWE-corrected). Cross-sectional images of the significant clusters as well as unthresholded statistical maps can be found in the Supplementary Information (Figure S7/S8). **d)** These clusters persisted when repeating the analyses while partialing out the effects of emotional word content (using affect grids), visual wordform (using a visual-categorization DNN), and auditory properties of the spoken words (using a speech-recognition DNN). **e)** Within the IPC cluster defined on the full 45-million sentence model (marked by an arrow), we compared model families trained on different corpus sizes and on only abstract or concrete words, respectively. Brain-like representations emerged in models that were trained on as little as 100,000 sentences and on either abstract or concrete embeddings. Dots show individual-participant data, error bars denote SEM, asterisks represent p<0.05 (FDR-corrected).

Cortical responses were modelled in a representational similarity analysis (RSA) framework. In RSA, the representational organization found in the brain is directly compared to the representational organization found in candidate representational models: this is done in a computationally straightforward way, whereby pairwise similarity relations are correlated across a larger set of stimuli (Kriegeskorte et al., 2008). Here, we used an RSA searchlight approach, in which we extracted similarity relations among the words across the whole cortex (Figure 1b; see Materials and Methods). We modelled these similarities using a word2vec model of distributional semantics (Mikolov et al., 2013), trained on a 45-million sentence corpus (SdeWaC; Faaß & Eckhart, 2013).

The model predicted brain activations in the left inferior parietal cortex (IPC; Figure 1c), most prominently covering the angular gyrus, but extending into the superior parietal and middle occipital cortices (158 voxels, peak: -36/-58/46, t[18]=6.16), and in right IPC, with an anterior cluster primarily in the supramarginal gyrus (91 voxels, peak: 45/-46/58, t[18]=4.91), and a posterior cluster in the angular gyrus (35 voxels, peak: 36/-73/40, t[18]=4.80). Detailed analyses, reported in the Supplementary Information, show that this correspondence was not driven by individual words present in the stimulus set (Figure S2). These results show that bilateral IPC represents semantic similarities of abstract concepts. They further suggest IPC as a cortical hub for the activation and contextualization of abstract concepts.

This interpretation, however, warrants a note of caution: Although it is tempting to strongly interpret the IPC representations as a reflection of concept coding, they may also reflect linguistic coding. While the distributional information extracted from word2vec inherently contains conceptual properties (e.g., Huebner & Willits, 2018; Tshitoyan et al., 2019; Utsumi et al., 2020), it is primarily linguistic. As also our experimental design used words to activate representations in the brain, the similarities between the model and the brain we see may be driven by the linguistic, rather than the conceptual information these words carry.

### Controlling for emotional and sensory word properties

Some researchers have argued that abstract concepts are represented through their grounding in the emotional domain (Kousta et al., 2011; Vigliocco et al., 2011). To test whether IPC representations are indeed driven by the words’ emotional content, we re-performed the analysis while partialing out valence and arousal ratings (see Materials and Methods). We still found clusters in left (36 voxels, peak: -33/-64/31, t[18]=4.73), and right (108 voxels, peak: 63/-40/31, t[18]=7.30) IPC, suggesting that the emotional content is insufficient to explain abstract concept representation in parietal cortex. Notably, the left IPC cluster somewhat shrunk after controlling for emotional word properties. This may be because word2vec models can pick up on emotional features implicitly contained in word embeddings (Utsumi, 2020).

IPC is also sensitive to sensory properties, such as visual form (Freud et al., 2018) and phonological speech attributes (Hartwigsen et al., 2010). However, when repeating the analysis while partialing out early activations in a visual-categorization deep neural network (DNN; see Materials and Methods), we still found clusters in the left (157 voxels, peak: -36/-58/46, t[18]=6.07) and right (anterior: 76 voxels, peak: 51/-49/52, t[18]=4.70; posterior: 32 voxels, peak: 39/-76/40, t[18]=4.82) IPC. Similarly, when controlling for activations obtained from a speech-recognition DNN (see Materials and Methods), we still found clusters in left (113 voxels, peak: -33/-64/31, t[18]=5.43, p_cluster_<0.001), and right (53 voxels, peak: 48/-46/55, t[18]=4.68) IPC. These results show that sensory properties are unrelated to IPC representations of abstract concepts.

In the Supplementary Information, we additionally show that word frequency cannot account for the correspondence between the word2vec model and brain representations in IPC (Figure S4).

### Trajectories towards brain-like representations

Our observation that the word2vec model and IPC share abstract concept representations led us to ask how the model acquires this property. To test whether co-occurrence statistics are acquired incrementally over increasing experience with human language, we devised a word2vec model family whose members were trained on staggered amounts of data, from the full 45-million sentence corpus down to fragments as small as 1,000 sentences. We then evaluated how well models trained on less data could still predict representations in the IPC cluster that yielded the best correspondence with the full 45-million sentence model (see Materials and Methods).

This analysis revealed decreasing correspondence with decreasing training data (mean r=0.74, t[18]=5.89, p<0.001). Nonetheless, a model trained on only 100,000 sentences (∼0.2% of the corpus) still predicted IPC representations well (t[18]=3.46, p_FDR_=0.002; comparison to full model: t[18]=2.15, p=0.045), whereas models trained on smaller training sets did not (Figure 1d). Direct comparisons of all models to the full model trained on 45m sentences can be found in the Supplementary Information (Figure S5). These results show that brain-like representations are learned through linguistic co-occurrence statistics, which can emerge already from a (relatively) modest degree of training experience.

### Modelling brain representations from abstract and concrete embeddings

Some theorists argue that the meaning of abstract concepts needs to be derived through the activation of related concrete concepts, which are in turn grounded in sensory experiences (Kiefer & Pulvermüller, 2012; Lakoff, 2008). This view prompts the hypothesis that representations of abstract concepts originate primarily from co-occurrence statistics between abstract and concrete words, rather than among abstract words alone. To test this hypothesis, we trained word2vec models on subsets of the 45-million sentence corpus that we devised to consist of abstract or concrete words only (see Materials and Methods).

Models trained on abstract-only and concrete-only corpora both predicted representations in IPC (Figure 1e). Reproducing the previous pattern of results, we found that models trained on larger fractions of the corpus predicted representations better (abstract only: mean r=0.36, t[18]=2.55, p=0.010; concrete-only: mean r=0.73, t[18]=4.32, p<0.001). Interestingly, representations were modelled equally well by the most extensively trained abstract-only (t[18]=3.09, p_FDR_=0.010) and concrete-only models (t[18]=2.60, p_FDR_=0.018; comparison: t[18]=0.50, p=0.62), suggesting that brain-like representations of abstract concepts can emerge from either abstract or concrete semantic embeddings.

## Conclusions

Our findings yield multiple key insights into abstract concept representation:

First, our findings provide novel evidence that the IPC is a core area for concept coding (Binder et al., 2009). Returning to the question of whether our results reflect genuine conceptual representation or language-specific codes, the localization of effects to the IPC indeed provides tentative evidence for the former: IPC activations are not routinely observed in language tasks (see Braga et al., 2020) – by contrast, particularly the angular gyrus is often implicated in brain networks for concept representation (Binder & Desai, 2011). Others, however, have recently contested this role of the IPC, as the region is not consistently activated during semantic cognition (Humphreys et al., 2015; Lambon Ralph et al., 2016). Critically, the current study used a task that required participants to activate and contextualize abstract concepts. In this task, we identify the angular gyrus as a critical hub for the dynamic use of abstract knowledge, consistent with the view that this region plays a key role in combinatory linguistic processing (David & Yee, 2019; Graessner et al., 2021; Price et al., 2015; Pylkkänen, 2019, 2020). Such combinatory processing may be particularly critical for abstract concepts, which more strongly need to be contextualized in a situational way during everyday use. It is worth noting, however, that our study does not establish that the angular gyrus is specifically important for representing abstract but not concrete words. In fact, previous results suggest that concrete words activate the angular gyrus just as strongly as abstract words (Hoffman et al., 2015), so that future studies need to carefully compare the representation of abstract and concrete concepts in this region.

Second, our study shows that brain representations of abstract concepts can be predicted from distributional word embeddings in natural language (Wang et al., 2018a). Interestingly, the organization of abstract concepts, as found in our brains, can be modelled from linguistic embeddings in both abstract and concrete realms of knowledge. This result shows that despite their representational dissimilarities (Binder et al., 2005; Wang et al., 2010), abstract and concrete concepts may be organized through shared principles. It is worth noting that the models constructed from abstract-only and concrete-only corpora in our study still produced a moderately high inter-correlation (r=0.78). Although this suggests a similarity in abstract words’ embeddings within other abstract and concrete words, this high correlation also limits the potential of our analysis to reveal substantial differences in how well these models predict neural representations. Future studies could specifically assemble stimulus sets that target concepts for which embeddings in the abstract and concrete realms are more different.

Third, our data informs theories of abstract knowledge representation (Borghi et al., 2017). Our results do not provide evidence for theories suggesting that abstract concepts are coded solely through emotional associations (Vigliocco et al., 2011) or the activation of related concrete concepts (Harpaintner et al., 2020; Lakoff, 2008). Further, in our study, we did not find evidence for an additional visual representation of abstract concepts (Anderson et al., 2017; Tang et al., 2021) or for a grounding of abstract conceptual knowledge in cognitive/motor systems (Dreyer & Pulvermüller, 2018). Our findings rather suggest that abstract knowledge is reflected in distributional relationships in neural representations of the concept or language processing systems. However, the question how distributional codes (such as the ones capitalized on by language models like word2vec) relate to word meaning is controversial: positions range from claims that word meaning is determined by distributed relations and respective neural codes akin to those in word2vec (Firth, 1957; Harris 1954; Landauer & Dumais, 1997) to the argument that distributional codes are insufficient to provide insights into meaning (Bender & Koller, 2020) – under this view, the observed similarities might be an emerging phenomenon rather than the underlying coding scheme. The current investigation cannot arbitrate between such theoretical positions.

Fourth, our results suggest that by harnessing co-occurrence statistics from linguistic experience, computational models of distributional semantics can acquire abstract concept representations that are organized in similar ways as biological representations. Although massive corpora are immensely popular for modelling language organization, our analyses of model learning trajectories show that brain-like representations can emerge from much smaller training sets of only 100,000 sentences. Cleary, our study provides only a first, coarse approximation of the tentative learning trajectory towards brain-like representations. Future work needs to map out the emergence of more fine-grained information along these learning trajectories to investigate how closely the acquisition of the models’ representations across training can predict human concept learning and development (Vigliocco et al., 2018).

Finally, our study highlights that computational models – through systematic manipulation of model training regimes – can yield targeted insights into the emergence and format of concept representations. Moving ahead, future studies could not only refine training regimes but also comprehensively manipulate a set of fundamental model parameters (Cichy & Kaiser, 2019). First advances have recently been made by comparing language models with different architectures to brain data (Schrimpf et al., 2021), by enriching models with predictive and contextual information (Anderson et al., 2021; Goldstein et al., 2020; Jain & Huth, 2018; Lopopolo et al., 2020) and by testing the applicability of linguistic models to experiences in domains like vision (Bonner & Epstein, 2021; Hayes & Henderson, 2021; van Paridon et al., 2021).In the future, employing such targeted model-based analyses may yield further fine-grained insights into how our brain represents abstract knowledge.

## Materials and Methods

### Participants

Nineteen healthy adults (mean age 28.8 years, SD=6.1; 10 female) with normal or corrected-to-normal vision completed the experiment. All of them were right-handed and native German speakers. Participants provided informed consent and received monetary reimbursement or course credits for participation. Participants were recruited from the online participant database of the Berlin School of Mind and Brain (Greiner, 2015). All procedures were approved by the local ethical committee and were in accordance with the Declaration of Helsinki.

### Stimuli and Paradigm

The stimulus set consisted of 61 abstract German nouns. These nouns were chosen from a list of the most frequent German words (from: wortschatz.uni-leipzig.de), from which they were arbitrary selected to cover a range of themes. All words and their English translations can be found in the Supplementary Information (Table S1).

During the fMRI experiment, participants completed 10 runs. Before each run, participants read through one of 10 contextual background stories. All texts and their English translations can be found in the Supplementary Information (Table S2). Participants were asked to mentally image themselves being in the scenario outlined in the text. After reading through the text, participants were instructed to use the subsequently presented words in the upcoming run to mentally narrate a story that incorporates the words as they are shown on the screen. They were instructed that it is completely up to them how the story unfolds as long as they use all the words in their story. Stories were chosen to be emotionally engaging to increase participants’ immersion into the task. The order of the 10 stories was randomized for every participant.

Each run contained 61 experimental trials. On each trial, one of the abstract words was shown for 3 seconds, in black Arial font on a gray background. Trials were separated by an inter-trial interval of 1.5 seconds, during which a fixation cross was shown. In addition to the experimental trials, each run included 14 fixation trials, where only the fixation cross was shown throughout the trial. Trial order was randomized within each run.

To ensure that participants paid attention to the words, we introduced a simple manual task: In each run, 7 of the word were shown in pink color and participants had to press a button whenever they saw one them.

Runs started and ended with brief fixation periods; each run lasted 5:48 minutes. The stimulation was back-projected onto a translucent screen at the end of the MRI scanner bore and controlled using the Psychtoolbox (Brainard, 1997).

Additionally, prior to the experiment, each participant completed a practice run (using a background text different from the ones used in the experiment).

Two participants completed a version of the experiment that differed in two aspects: the inter-trial interval was 1s instead of 1.5s and no behavioral task was included.

### MRI acquisition and preprocessing

MRI data was acquired using a 3T Siemens Tim Trio Scanner equipped with a 12-channel head coil. T2*-weighted gradient-echo echo-planar images were collected as functional volumes (TR=2s, TE=30ms, 70° flip angle, 3mm^3^ voxel size, 37 slices, 20% gap, 192mm FOV, 64×64 matrix size, interleaved acquisition). Additionally, a T1-weighted image (MPRAGE; 1mm^3^ voxel size) was obtained as a high-resolution anatomical reference. Preprocessing was done in MATLAB using SPM12 (www.fil.ion.ucl.ac.uk/spm/). The functional volumes were realigned and coregistered to the T1 image. The T1 image was normalized to MNI-305 standard space to obtain transformation parameters used to normalize participant-specific results maps (see below).

### Representational similarity analysis

To quantify neural representations, we used multivariate representational similarity analysis (RSA) (Kriegeskorte et al., 2008). In RSA, neural representations are first characterized by means of their pairwise similarity structure (i.e., how similarly each stimulus is represented with each other stimulus). The pairwise dissimilarities between neural representations are organized in neural representational dissimilarity matrices (RDMs) indexed in rows and columns by the experimental conditions compared. Then, the neural similarity structure (i.e., the neural RDMs) are correlated to model RDMs, which capture different aspects of the conditions’ similarity. Significant correlations between the neural RDMs and these model RDMs indicate that the aspect of similarity conveyed by the model is represented in the brain.

In recent studies, RSA has successfully been used to relate representations in computational models of language processing and human cortex (Lopopolo et al., 2020; Martin et al., 2018; Wang et al., 2018a, 2018b). Other studies have probed correspondences between language processing models and the brain through the use of encoding models (Bonner & Epstein, 2021; Deniz et al., 2019; Huth et al., 2016; Jain & Huth, 2020; Pereira et al., 2018). Encoding models seek to directly establish a mapping between the feature dimensions extracted by the computational model and fMRI responses in individual voxels. For this task, they require diverse and large sets of data to train the model weights (van Gerven, 2017), which the current design was not optimized for. RSA and encoding models offer largely complimentary quantifications of neural representations, with comparable sensitivity (Diedrichsen & Kriegeskorte, 2017) and comparable constraints regarding interpretability (Kriegeskorte & Douglas, 2019). However, although RSA offers a straightforward and computationally efficient way to relate computational models and population codes in the brain, one limitation needs to be taken into account when interpreting the results: representations in some parts of the brain may rely on intricate weightings of few feature dimensions and may therefore be harder to identify with RSA than with encoding models.

#### Extracting neural dissimilarity

Separately for each participant and each run, we first modeled the functional MRI data in a general linear model (GLM) with 67 predictors (61 predictors for the 61 words, and 6 predictors for the 6 movement regressors obtained during realignment). From these GLMs, we obtained 610 beta weights of interest for every voxel, which quantified the voxel’s activation to each of the 61 words in each of the 10 runs. All further analyses were carried out using a searchlight approach (Kriegeskorte et al., 2006), that is, analyses were done repeatedly for a spherical neighborhood (3-voxel radius) centered on each voxel across the brain. This approach allowed us to quantify and model neural representations in a continuous and unconstrained way across brain space.

For each searchlight neighborhood, neural RDMs were created based on the similarity of multi-voxel response patterns, using the CoSMoMVPA toolbox (Oosterhof et al., 2016). Within each neighborhood, we extracted the response pattern across voxels evoked by each word in each run. We then performed a cross-validated correlation analysis (Haxby et al., 2001). Unbiased, cross-validated distance metrics like the cross-validated correlations are generally considered more reliable than non-cross-validated metrics (such as plain correlations) for estimating pattern similarities in brain data (Walther et al., 2016). For this analysis, the data were repeatedly split into two halves (all possible 50/50 splits; results were later averaged across these splits) and the response patterns for each word were averaged within each half. For each pair of words, we then computed two correlations: (i) within-condition correlations were computed by correlating the response patterns evoked by each of the two words in one half of the data with the response patterns evoked by the same word in the other half of the data, and (ii) between-condition correlations were computed by correlating the response patterns evoked by each of the two words in one half of the data with the response patterns evoked by the other word in the other half of the data. By subtracting the between-correlations from the within-correlations for each pair of words, we obtained an index of how dissimilar two words are based on the response patterns they evoked in the current searchlight neighborhood. Repeating this analysis for each pair of words yielded a 61×61 neural RDM for each searchlight.

#### Modelling neural dissimilarity

To model the semantic representation of the abstract words, we used a word2vec computational model of distributional semantics (Mikolov et al., 2013). The model was trained on the SdeWaC corpus, which contains 45 million German sentences (Faaß & Eckhart, 2013), using the gensim library (https://github.com/RaRe-Technologies/gensim). The model hyperparameters were the following: dimensions = 300, model type = skipgram, windowsize = 5, minimum count = 1, iterations = 50. For each word in the corpus, this model yields a vector representation that indicates its position in a 300-dimensional vector space. Distances in this vector space reflect similarities in word embeddings. We then created a 61×61 RDM based on the pairwise correlations of the 300 vector-space features for each of the words used in the experiment.

To establish correspondences between the model and the brain data, the model RDMs were correlated with the neural RDMs for each searchlight. These correlations were then Fisher-transformed and mapped back to the searchlight center. We thereby obtained brain maps of correspondence between each model and the neural data. For each participant, these maps were warped into standard space by using the normalization parameters obtained during preprocessing.

#### Controlling for emotional, visual, and auditory word properties

As an emotional content model, we used participants’ responses in an affect grid task, where 20 participants (partly including the participants in the current experiment) concurrently rated each word’s valence and arousal by selecting one compartment of a 9×9 grid (Russell et al., 1989). From these data, we created two RDMs: (i) a valence RDM, whose entries reflected pairwise absolute difference in the words’ valence ratings and (ii) an arousal RDM, whose entries reflected pairwise absolute difference in the words’ arousal ratings. The valence and arousal RDMs were mildly correlated with each other (r=0.19) and with the different word2vec model RDMs (all r<0.24). The words’ similarity in valence an arousal did not significantly predict brain activations in a searchlight analysis.

As a visual word form model, we used activations in the three earliest convolutional layers an AlexNet DNN pre-trained on object recognition (Krizhevsky et al., 2012; Vedaldi & Lenc, 2015), which have been shown to capture representations of simple visual attributes in visual cortex (Cichy et al., 2016). We printed the 61 words as they appeared in the experiment on a 225×225 pixel gray image background and fed the resulting images to the background. The resulting network activations were used to construct model RDMs. For each of the first three convolutional layers of the network, the RDM was constructed by computing pairwise distances (1-correlation) between layer-specific activation vectors. The visual DNN RDMs were only very weakly correlated with the word2vec model RDMs (all r<0.1). Searchlight analyses revealed that the first three layers of the visual DNN predicted activations in bilateral posterior visual cortex, including fusiform cortex (see Supplementary Information, Figure S1).

As a model of auditory, phonetic word similarity, we used activations in a DNN model of auditory speech recognition (Kell et al., 2018). We obtained spoken versions of the 61 words from the ttsmp3 webpage (https://ttsmp3.com/text-to-speech/German/). The sound files were resized to a length of 2 seconds by right-padding them with zeros, transformed into a cochleagram representation, and then passed through the speech recognition branch of the DNN. The resulting network activations were used to construct model RDMs. For each of the seven layers of the network, the RDM was constructed by computing pairwise distances (1-correlation) between layer-specific activation vectors. The auditory DNN RDMs were only weakly correlated with the word2vec model RDMs (all r<0.16). Searchlight analyses revealed that the early layers of the auditory DNN, because of the correlation between word length and speech duration, also predicted activations in bilateral posterior visual cortex. By contrast, the last layer of the network specifically predicted activations in left middle temporal gyrus (see Supplementary Information, Figure S1).

To control for emotional and sensory properties, we performed searchlight analyses relating the neural RDMs and the word2vec model RDMs as before, while we partialed out the two emotion predictor RDMs, the three visual DNN predictor RDMs, or the seven auditory DNN predictor RDMs, respectively. Specifically, for each searchlight neighborhood, we computed a partial correlation between the neural RDM and the predictor RDM which was controlled for the control RDMs. This procedure ensured that if the control RDMs predicted the same portion of variance in the neural RDM as the predictor RDM, the correlation would disappear (for similar approaches, see Ambrus et al., 2019; Cichy et al., 2019; Kaiser & Nyga, 2020). All other aspects of the analysis remained identical to the previous searchlight analysis.

#### Region of interest analyses

For further dissecting the representations in left parietal cortex, we specifically focused on this area in a region-of-interest (ROI) analysis. The IPC clusters that showed significant correspondence with the word2vec model in the main analysis were chosen as the ROI. Neural RDMs were generated from pairwise correlations of activity patterns across all voxels in the ROI; the procedure was otherwise identical to the procedure applied in the searchlight analysis (see above).

As ROI definition was done on the basis of the model that was trained on the full 45-million sentences SDeWaC corpus, we never evaluated this model statistically in our ROI analysis. We instead probed the correspondence between neural RDMs in the ROI and RDMs built from a set of different word2vec model families whose training regimes differed in important aspects.

To probe the behavior of our word2vec model with changes in training set, we created a model family whose members were trained on different amounts of data. Models were trained on different fragments of the corpus (containing 45m, 10m, 1m, 100k, 10k, or 1k sentences). Each of these fragments corresponded to the first *n* sentences in the corpus (e.g., the 1k model comprised the first 1,000 sentences). We thereby ensured that the smaller fragments were always completely included in the larger ones.

Additionally, we constructed a model family whose members were trained on abstract words and a model family whose members were trained on concrete words. Members in each family differed by the amount of data they were trained on (as outlined above). Abstract and concrete words were defined on the basis of the abstractness-concreteness scale of the IMS norms (https://www.ims.uni-stuttgart.de/en/research/resources/experiment-data/affective-norms) (Köper & im Walde, 2017). For the abstract-only models, we chose words that had a z-value of <0 on the on this scale and removed all other words from the corpus; this left us with ∼65 million words (∼4% of the corpus). For the concrete-only models, we chose words that had a z-value of >0 and removed all other words from the corpus; this left us with ∼280 million words (∼18% of the corpus). Note that for the concrete-only model, the 61 abstract words were also left in the corpus, so that relationships between them and the concrete words could be obtained.

For all models of each model family, we extracted a 61×61 RDM, which was then correlated with the neural RDM extracted for the ROI; these correlations were Fisher-transformed before statistical analysis. Correlations between all RDMs constructed from the word2vec models can be found in the Supplementary Information (Figure S3).

### Statistical testing

For the searchlight analyses, to detect spatial clusters in which the neural data were explained by the different representational models, we performed one-sided t-tests against zero across participants, separately for each voxel in the correlation maps. The resulting statistical maps were thresholded at the voxel level at p_voxel_<0.001 (uncorrected) and at the cluster level at p_cluster_<0.05 (family-wise error corrected, as implemented in SPM12). These thresholds were selected based on recommendations for cluster-based thresholding of fMRI results (Woo et al., 2014). In the Supplementary Information (Figure S6), we show that similar results are reached with an alternative statistical approach based on threshold-free cluster enhancement (TFCE; Smith & Nichols, 2009).

For the cross-validated ROI analyses, ROIs were repeatedly defined in 18 participants, using the same statistical thresholding as in the full searchlight analysis (p_voxel_<0.001, p_cluster_<0.05 FWE-corrected). Correlations between neural RDMs extracted from the ROIs and model RDMs were then evaluated using one-sided t-tests against zero across participants. Results were corrected for multiple comparisons across the different training corpus sizes using FDR corrections.

## Data availability

Data are publicly available on OSF (https://doi.org/10.17605/OSF.IO/FTBJQ). For other materials, please contact the corresponding author.

## Acknowledgements

Thanks to Raphael Leuner and Kshitij Dwivedi for help with the computational analyses. D.K. and R.M.C. are supported by DFG grants (KA4683/2-1, CI241/1-1, CI241/3-1, CI241/7-1). R.M.C. is supported by an ERC Starting Grant (ERC-2018-StG 803370).

## References

Ambrus GG, Kaiser D, Cichy RM, Kovács G. (2019). The neural dynamics of familiar face recognition. Cereb Cortex, 29, 4775–4784.

Anderson AJ, et al. (2021). Deep artificial neural networks reveal a distributed cortical network encoding propositional sentence-level meaning. J Neurosci, 41, 4100–4119.

Anderson AJ, Kiela D, Clark S, Poesio M. (2017). Visually grounded and textual semantic models differentially decode brain activity associated with concrete and abstract nouns. Trans Assoc Comput Linguist, 5, 17–30.

Anderson AJ, Murphy B, Poesio M. (2014). Discriminating taxonomic categories and domains in mental simulations of concepts of varying concreteness. J Cogn Neurosci, 26, 658–681.

Bender EM, Koller A. (2020). Climbing towards NLU: on meaning, form, and understanding in the age of data. Proceedings of the 58^th^ Annual Meeting of the Association for Computational Linguistics, 5185–5198.

Binder JR, Desai RH. (2011). The neurobiology of semantic memory. Trends Cogn Sci, 15, 527–536.

Binder JR, Desai RH, Graves WW, Conant LL. (2009). Where is the semantic system? A critical review and meta-analysis of 120 functional neuroimaging studies. Cereb Cortex, 19, 2767–2797.

Binder JR, Westbury CF, McKiernan KA, Possing ET, Medler DA. (2005). Distinct brain systems for processing concrete and abstract concepts. J Cogn Neurosci, 17, 905–917.

Bonner MF, Epstein RA. (2021). Object representations in the human brain reflect the co-occurrence statistics of vision and language. Nat Commun, 12, 4081.

Borghi AM, et al. (2017). The challenge of abstract concepts. Psychol Bull, 143, 263–292.

Braga RM, DiNicola LM, Becker HC, Buckner RL. (2020). Situating the left-lateralized language network in the broader organization of multiple specialized large-scale distributed networks. J Neurophysiol, 124, 1415–1448.

Brainard DH. (1997). The psychophysics toolbox. Spat Vis, 10, 433–436.

Cichy RM, Kaiser D. (2019). Deep neural networks as scientific models. Trends Cogn Sci, 23, 305–317.

Cichy RM, Khosla A, Pantazis D, Torralba A, Oliva A. (2016). Comparison of deep neural networks to spatio-temporal cortical dynamics of human visual object recognition reveals hierarchical correspondence. Sci Rep, 6, 27755.

Cichy RM, Kriegeskorte N, Jozwik KM, van den Bosch JJF, Charest I. (2019). The spatiotemporal neural dynamics underlying perceived similarity for real-world objects. Neuroimage, 194, 12–24.

Deniz F, Nunez-Elizalde AO, Huth AG, Gallant JL. (2019). The representation of semantic information across human cerebral cortex during listening versus reading is invariant to stimulus modality. J Neurosci, 39, 7722–7736.

Diedrichsen J, Kriegeskorte N. (2017). Representational models: a common framework for understanding encoding, pattern-component, and representational-similarity analysis. Plos Comput Biol, 13, e1005508.

Dreyer FR, Pulvermüller, F. (2018). Abstract semantics in the motor system? – An event-related fMRI study on passive reading of semantic word categories carrying abstract emotional and mental meaning. Cortex, 100, 52–70.

Faaß G, Eckhart K. (2013). SdeWaC – a corpus of parsable sentences from the web. In: Gurevych I, Biemann C, Zesch T (eds). Language processing and knowledge in the web. Lecture notes in computer science, vol. 8105. Springer, Berlin, Heidelberg.

Firth JR. (1957). A synopsis of linguistic theory, 1930-1950. In: Studies in Linguistic Analysis. Blackwell, Oxford, UK.

Freud E, Culham JC, Plaut DC, Behrmann M. (2018). The large-scale organization of shape processing in the ventral and dorsal pathways. eLife, 6, e27576.

Goldstein A, et al. (2021). Thinking ahead: spontaneous prediction in context as a keystone of language in humans and machines. bioRxiv, doi.org/10.1101/2020.12.02.403477.

Graessner A, Zaccarella E, Hartwigsen G. (2021). Differential contributions of left-hemispheric language regions to basic semantic composition. Brain Struct Funct, 226, 501–518.

Greiner B. (2015). Subject pool recruitment procedures: organizing experiments with ORSEE. JESA, 1, 114–125.

Hartwigsen G, et al. (2010). Phonological decisions require both the left and right supramarginal gyri. Proc Natl Acad Sci USA, 107, 16494–16499.

Harpaintner M, Sim E-J, Trumpp NM, Ulrich M, Kiefer M. (2020). The grounding of abstract concepts in the motor and visual system: an fMRI study. Cortex, 124, 1–22.

Harris ZS. (1954). Distributional structure. Word, 10, 146–162.

Haxby JV, et al. (2001). Distributed and overlapping representations of faces and objects in ventral temporal cortex. Science, 293, 2425–2430.

Hayes TR, Henderson JM. (2021). Looking for semantic similarity: what a vector-space model of semantics can tell us about attention in real-world scenes. Psychol Sci, doi.org/10.1177/0956797621994768.

Hoffman P, Binney RJ, Lambon Ralph MA. (2015). Differing contributions of inferior prefrontal and anterior temporal cortex to concrete and abstract conceptual knowledge. Cortex, 63, 250–266.

Huebner PA, Willits JA. (2018). Structured semantic knowledge can emerge automatically from predicting words sequences in child-directed speech. Front Psychol, 9, 133.

Humphreys GF, Hoffman P, Visser M, Binney RJ, Lambon Ralph MA. (2015). Establishing task-and modality-dependent dissociations between the semantic and default mode networks. Proc Natl Acad Sci USA, 112, 7857–7862.

Huth AG, de Heer WA, Griffiths TL, Theunissen FE, Gallant JL. (2016). Human natural speech reveals the semantic maps that tile human cerebral cortex. Nature, 532, 453–458.

Jain S, Huth AG. (2018). Incorporating context into language encoding models for fMRI. Advances in Neural Information Processing Systems, 31.

Just MA, Cherkassky VL, Aryal S, Mitchell TM. (2010). A neurosemantic theory of concrete noun representation based on the underlying brain codes. PLoS One, 5, e8622.

Kaiser D, Nyga K. (2020). Tracking cortical representations of facial attractiveness using time-resolved representational similarity analysis. Sci Rep, 10, 16852.

Kell AJ, Yamins DL, Shook EN, Norman-Haignere SV, McDermott JH. (2018). A task-optimized neural network replicates human auditory behavior, predicts brain responses, and reveals a cortical processing hierarchy. Neuron, 98, 630–644.

Kiefer M, Pulvermüller F. (2012). Conceptual representations in mind and brain: Theoretical developments, current evidence and future directions. Cortex, 48, 805–825.

Köper M, im Walde SS. (2017). Improving verb metaphor detection by propagating abstractness to words, phrases and individual senses. Proceedings of the 1st workshop on sense, concept and entity representations and their applications, 24–30.

Kousta ST, Vigliocco G, Vinson DP, Andrews M, Del Campo E. (2011). The representation of abstract words: why emotion matters. J Exp Psychol Gen, 140, 14.

Kriegeskorte N, Douglas PK. (2019). Interpreting encoding and decoding models. Curr Opin Neurobiol, 55, 167–179.

Kriegeskorte N, Goebel R, Bandettini P. (2006). Information-based functional brain mapping. Proc Natl Acad Sci USA, 103, 3863–3868.

Kriegeskorte N, Mur M, Bandettini P. (2008). Representational similarity analysis – connecting the branches of systems neuroscience. Front Syst Neurosci, 2, 4.

Krizhevsky A, Sutskever I, Hinton GE. (2012). ImageNet classification with deep convolutional neural networks. Advances in Neural Information Processing Systems, 1097–1105.

Lakoff G. (2008). The neural theory of metaphor. In: Gibbs Jr, WR (ed). The Cambridge handbook of metaphor and thought. Cambridge University Press, Cambridge.

Landauer TK, Dumais ST. (1997). A solution to Plato’s problem: the latent semantic analysis theory of acquisition, induction, and representation of knowledge. Psychol Rev, 104, 211–240.

Lopopolo A, Schoffelen JM, van den Bosch A, Willems RM. (2020). Words in context: tracking context-processing during language comprehension using computational language models and MEG. bioRxiv, doi.org/10.1101/2020.06.19.161190.

Martin CB, Douglas D, Newsome RN, Man LLY, Barense MD. (2018). Integrative and distinctive coding of visual and conceptual object features in the ventral visual stream. eLife, 7, e31873.

Mikolov T, Chen K, Corrado G, Dean J. (2013). Efficient estimation of word representations in vector space. arXiv, 1301.3781.

Mitchell TM, et al. (2008). Predicting human brain activity associated with the meanings of nouns. Science, 320, 1191–1195.

Oosterhof NN, Connolly AC, Haxby JV. (2016). CoSMoMVPA: Multi-modal multivariate pattern analysis of neuroimaging data in Matlab/GNU Octave. Front Neuroinform, 10, 20.

Pereira F, et al. (2018). Toward a universal decoder of linguistic meaning from brain activation. Nat Commun, 9, 1–13.

Price AR, Bonner MF, Peelle JE, Grossman M. (2015). Converging evidence for the neuroanatomic basis of combinatorial semantics in the angular gyrus. J Neurosci, 35, 3276–3284.

Pylkkänen L. (2019). The neural basis of combinatory syntax and semantics. Science, 355, 62–66.

Pylkkänen L. (2020). Neural basis of basic composition: what we have learned from the red-boat studies and their extensions. Phil Trans R Soc Lond B Biol Sci, 375, 20190299.

Russell JA, Weiss A, Mendelsohn GA. (1989). Affect grid: a single-item scale of pleasure and arousal. J Personal Soc Psychol, 57, 493.

Schrimpf M, et al. (2021). The neural architecture of language: integrative modeling converges on predictive processing. bioRxiv, doi.org/10.1101/2020.06.26.174482.

Smith SM, Nichols TE. (2009). Threshold-free cluster enhancement: addressing problems of smoothing, threshold dependence and localization in cluster inference. Neuroimage, 44, 83–98.

Tang J, LeBel A, Huth AG. (2021). Cortical representations of concrete and abstract concepts in language combine visual and linguistic representations. bioRxiv, doi.org/10.1101/2021.05.19.444701.

Tshitoyan V, et al. (2019). Unsupervised word embeddings capture latent knowledge from materials science literature. Nature, 571, 95–98.

Utsumi A. (2020). Exploring what is encoded in distributional word vectors: a neurobiologically motivated analysis. Cogn Sci, 44, e1284.

van Gerven MAJ. (2017). A primer on encoding models in sensory neuroscience. J Math Psychol, 76, 172–183.

van Paridon J, Liu Q, Lupyan G. (2021). How do blind people know that blue is cold? Distributional semantics encode color-adjective associations. OSF preprints, doi.org/10.31234/osf.io/vyxpq

Vargas R, Just MA. (2020). Neural representations of abstract concepts: identifying underlying neurosemantic dimensions. Cereb Cortex, 30, 2157–2166.

Vedaldi A, Lenc K. (2015). MatConvNet – convolutional neural networks for Matlab. ACM International Conference on Multimedia.

Vigliocco G, et al. (2011). The neural representation of abstract words: the role of emotion. Cereb Cortex, 24, 1767–1777.

Vigliocco G, Ponari M, Norbury C. (2018). Learning and processing abstract words and concepts: insights from typical and atypical development. Top Cogn Sci, 10, 533–549.

Walther A., et al. (2016). Reliability of dissimilarity measures for multi-voxel pattern analysis. Neuroimage, 137, 188–200.

Wang J, Conder JA, Blitzer DN, Shinkareva SV. (2010). Neural representation of abstract and concrete concepts: a meta-analysis of neuroimaging studies. Hum Brain Mapp, 31, 1459–1468.

Wang X, et al. (2018a). Organizational principles of abstract words in the human brain. Cereb Cortex, 28, 4305–4318.

Wang X, et al. (2018b). Representational similarity analysis reveals task-dependent semantic influence on the visual word form area. Sci Rep, 8, 3047.

Woo C-W, Krishnan A, Wager TD. (2014). Cluster-extent based thresholding in fMRI analyses: pitfalls and recommendations. Neuroimage, 91, 412–419.

