## Supplementary Information for "Modelling brain representations of abstract concepts"

Supplementary Contents | Page

**Figure S1. Neural correlates of visual and auditory word similarity.** 2
**Figure S2. Individual-word effects.** 3
**Figure S3. Model intercorrelations.** 4
**Figure S4. Controlling for word frequencies.** 5
**Figure S5. Direct model comparisons in left IPC.** 6
**Figure S6. Whole-brain searchlight with TFCE statistics.** 7
**Figure S7. Searchlight results on cross-sectional images.** 8
**Figure S8. Unthresholded searchlight results.** 9

**Table S1. Abstract words.** 10
**Table S2. Background contexts.** 11

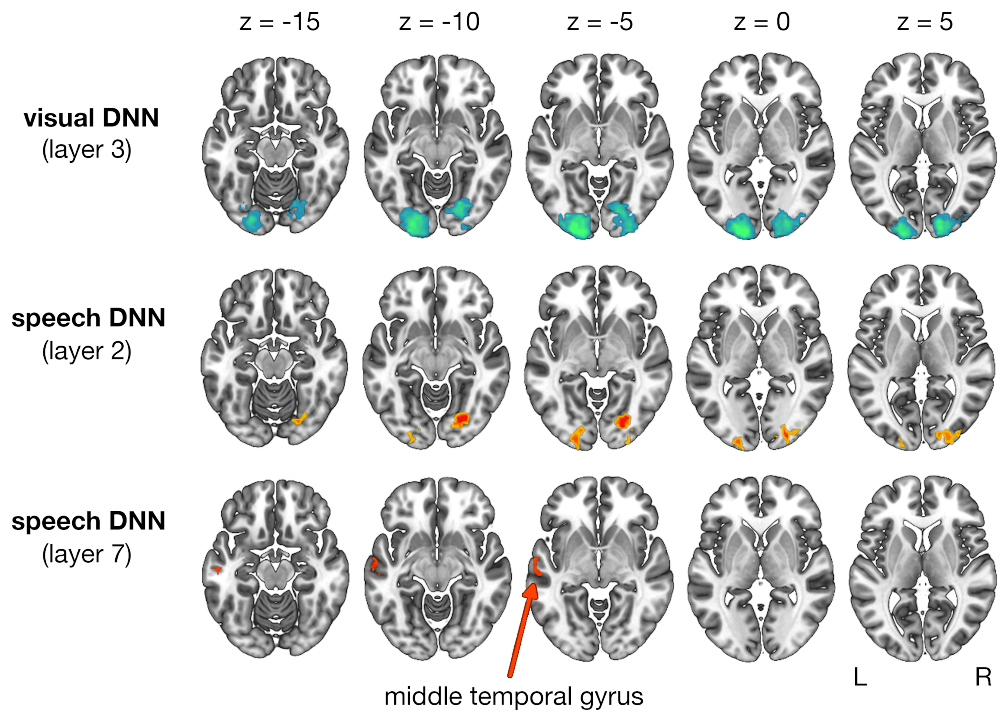


**Figure S1. Neural correlates of visual and auditory word similarity.** Visual word similarity, as modelled by the early layers of the an AlexNet DNN (here: layer 3) predicted activations in bilateral posterior visual cortex, including fusiform cortex (866 voxels, peak: -24/-97/-5, t[18]=7.89). Auditory similarity of the spoken words was modelled by a speech recognition DNN. Early layers of the network (here: layer 2), because of the correlation of word length and speech duration, also predicted activations in bilateral posterior visual cortex (266 voxels, peak: 24/-76/-8, t[18]=6.51). By contrast, the last layer of the network (layer 7) predicted activations in left middle temporal gyrus (34 voxels, peak: -60/-13/-8, t[18]=5.61). Brain maps are thresholded at $p_{voxel}<0.001$ (uncorrected) and $p_{cluster}<0.05$ (FWE-corrected).

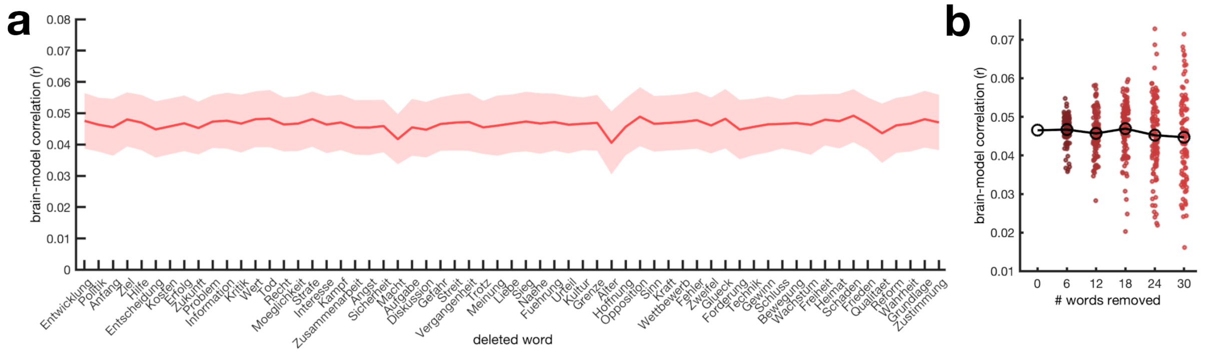


**Figure S2. Individual-word effects. a)** Brain-model correlations in left IPC when individual words were deleted from the model RDMs and neural RDMs before performing the analysis. The relatively homogeneous pattern shows that no single word exerted a substantial influence on the correspondence between model and brain. Error margins denote standard errors of the mean. **b)** Brain-model correlations in left IPC when removing a subset of up to 30 words at random from the RDMs. Results across 100 analyses with random subsets removed reveal that the pattern largely holds for smaller subsets of the stimulus space. All data points are means across all participants.

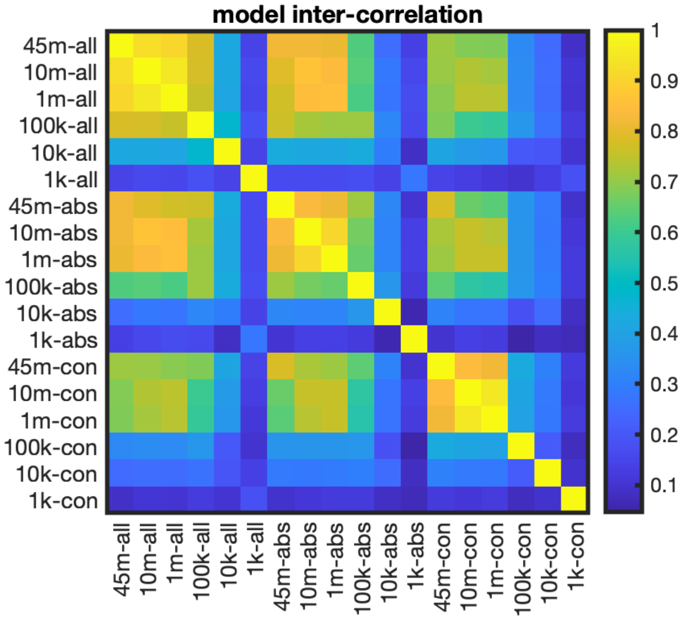


**Figure S3. Model intercorrelations.** Pairwise correlations between the representational dissimilarity matrices (RDMs) constructed for all word2vec model variants used in the study.

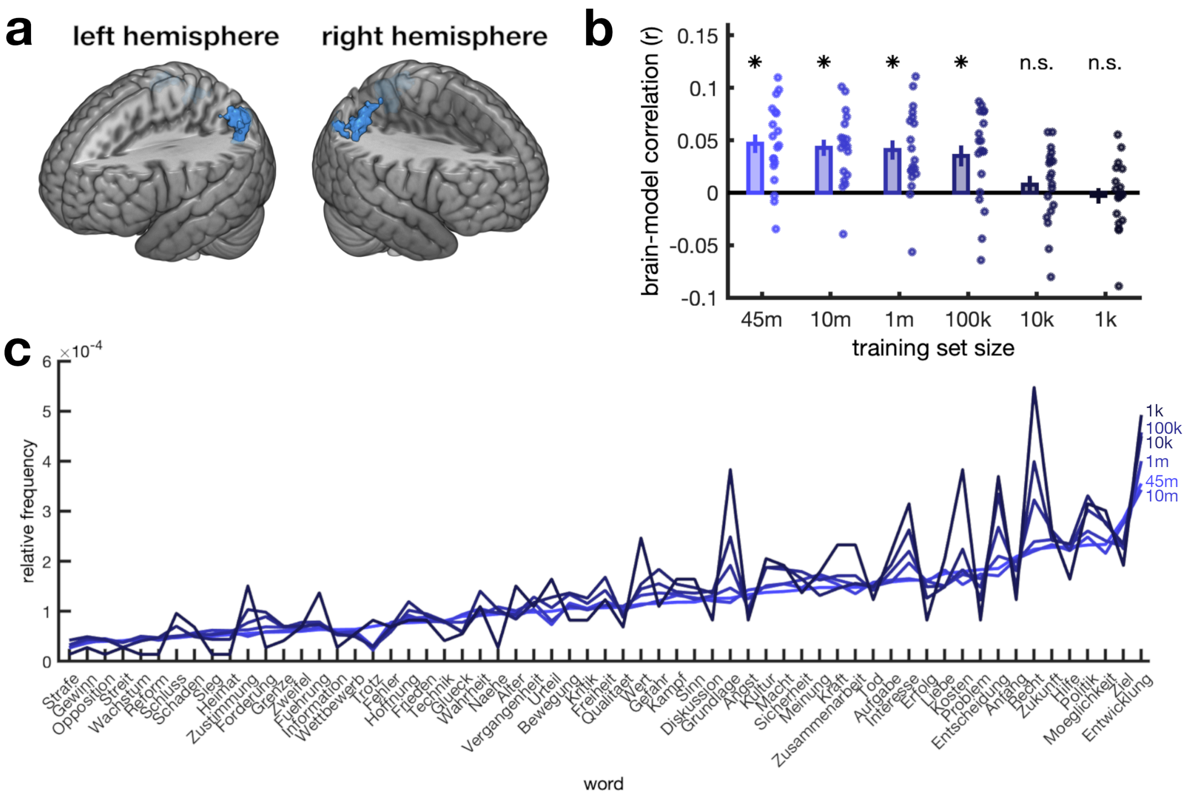


**Figure S4. Controlling for word frequencies. a)** Whole-brain searchlight analysis when controlling for similarities in word frequency using partial correlations. This analysis yielded results analogous to the original analysis (Figure 1c), with two clusters in right IPC (80 voxels, peak: 45/-46/58, t[18]=4.91, and 35 voxels, peak: 36/-73/40, t[18]=4.79) and one cluster in left IPC (163 voxels, peak: -36/-58/46, t[18]=6.23). **b)** Region-of-interest analysis in IPC for differently sized training corpora when similarities in word frequencies were controlled for. This analysis revealed essentially identical results to the main analysis (Figure 1e). **c)** Relative frequencies of the 61 abstract words across the differently sized training corpora. Despite some variations across corpus size, the words that were frequent in large corpora were also more frequent in the small corpora. Words are sorted by frequency in the largest (45m) corpus.

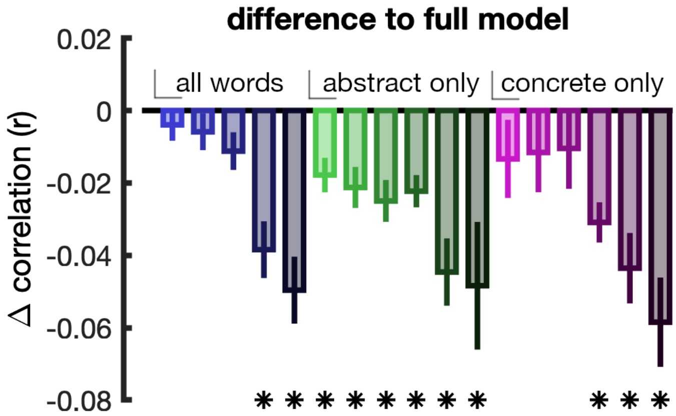


**Figure S5. Direct model comparisons in left IPC.** Differences between in brain-model correlations between the full model (45m sentences and all words included) and the trimmed models. Asterisks indicate significant differences to the full model, FDR-corrected for multiple comparisons. The full model itself is omitted from the plot. Color conventions are the same as in Figure 1e. Error bars denote standard errors.

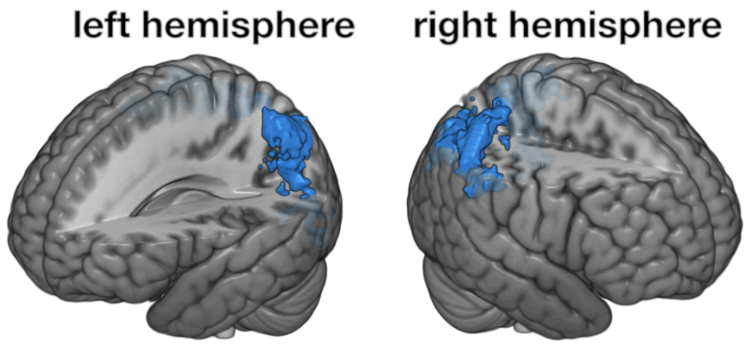


**Figure S6. Whole-brain searchlight with TFCE statistics.** Here, we used an alternative statistical test, based on threshold-free cluster enhancement (TFCE; Smith & Nichols, 2009), as implemented in CoSMoMVPA (Oosterhof et al., 2016). Z-scores for TFCE values were obtained by comparing the actual values to values across a null distribution constructed from 10,000 sign permutations. The resulting statistical maps were thresholded at $z>1.96$ ($p<0.05$). As in the main analysis (Figure 1c), two clusters emerged in right parietal cortex (503 voxels, peak: 42/-46/58, z=2.58, and 88 voxels, peak: -6/-58/58, z=2.11) and one cluster emerged in left parietal cortex (442 voxels, peak: -27/-73/46, z=2.89). Although also centered on the IPC, TFCE yielded somewhat more liberal results, with clusters extending more into the superior parietal cortices. No other clusters emerged across the brain.

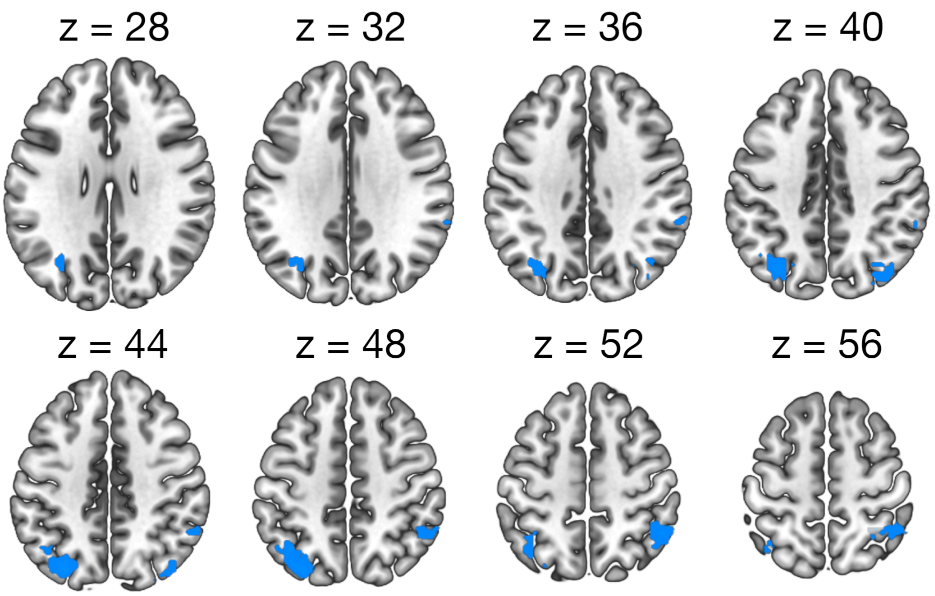


**Figure S7. Searchlight results on cross-sectional images.** Coronal brain slices are overlaid with regions that show significant correlations between neural representations and the full word2vec model (as in Figure 1c).

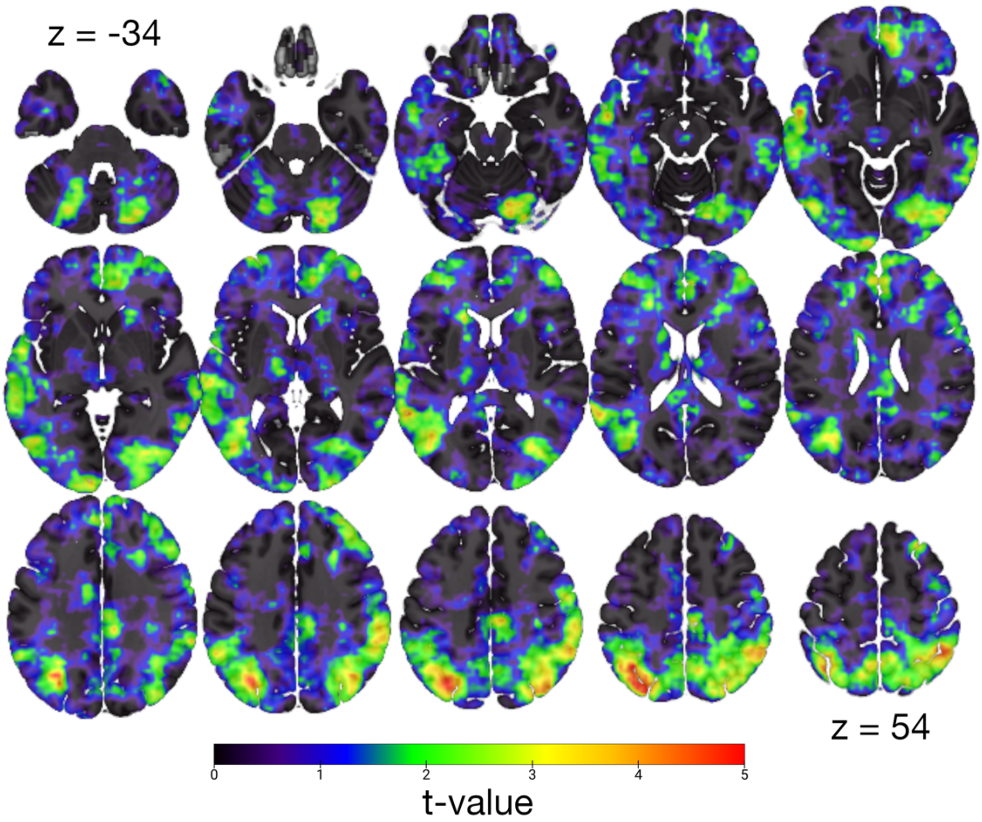


**Figure S8. Unthresholded searchlight results.** Coronal brain slices are overlaid with unthresholded t-maps comparing brain-model correlations to zero for the full word2vec model (as in Figure 1c). Slices are spaced continuously between z=-34 and z=54. Negative t-values are truncated.

**Table S1. Abstract words.** The 61 abstract German words used in the experiment and their English translations.

| German Word | English Translation | German Word | English Translation |
|---|---|---|---|
| Entwicklung | *development* | Recht | *law* |
| Politik | *politics* | Möglichkeit | *possibility* |
| Anfang | *beginning* | Strafe | *punishment* |
| Ziel | *goal* | Interesse | *interest* |
| Hilfe | *help* | Kampf | *fight* |
| Entscheidung | *decision* | Zusammenarbeit | *collaboration* |
| Kosten | *cost* | Angst | *anxiety* |
| Erfolg | *success* | Sicherheit | *safety* |
| Zukunft | *future* | Macht | *might* |
| Problem | *problem* | Aufgabe | *task* |
| Information | *information* | Diskussion | *discussion* |
| Zweifel | *doubt* | Gefahr | *danger* |
| Glück | *fortune* | Streit | *argument* |
| Forderung | *demand* | Vergangenheit | *past* |
| Technik | *technique* | Freiheit | *freedom* |
| Gewinn | *profit* | Heimat | *home* |
| Schluss | *end* | Schaden | *damage* |
| Bewegung | *movement* | Wettbewerb | *competition* |
| Wachstum | *growth* | Qualität | *quality* |
| Frieden | *peace* | Reform | *reform* |
| Fehler | *error* | Wahrheit | *truth* |
| Kritik | *critique* | Grundlage | *foundation* |
| Wert | *value* | Zustimmung | *agreement* |
| Tod | *death* | Urteil | *verdict* |
| Trotz | *defiance* | Kultur | *culture* |
| Meinung | *opinion* | Grenze | *border* |
| Liebe | *love* | Alter | *age* |
| Sieg | *victory* | Hoffnung | *hope* |
| Nähe | *closeness* | Opposition | *opposition* |
| Führung | *leadership* | Sinn | *sense* |
| Kraft | *force* | | |

**Table S2. Background contexts.** The 10 background contexts used in the experiment and their English translations. The gray text on the bottom was always used during the practice run.

| German Background Text | English Translation |
|---|---|
| Stell Dir vor Du bist ein Astronaut und explorierst ferne Galaxien. Dort triffst Du auf Angehörige einer bislang unbekannten Alienspezies. Versuche ihnen die Menschheit und ihre Kultur zu erklären. | *Imagine you are an astronaut exploring far-away galaxies. There you meet members of a previously unknown alien race. Try to explain humanity and its culture to them.* |
| Stell Dir vor Du bist Trauzeuge auf der Hochzeit Deines besten Freundes. Auf der Feier hältst Du eine Rede, in der Du einige gemeinsam durchlebte Anekdoten Revue passieren lässt. | *Imagine you are the best man / maid of honor at your best friend's wedding. At the party, you are giving a speech, in which you recall the past and tell a few anecdotes from your common past.* |
| Stell Dir vor Du bist ein Abenteurer auf dem Weg zum Nordpol. Du schreibst ein Kapitel Deines Reiseberichts in Dein Tagebuch, und betonst, trotz schwindender Vorräte und großer Kälte, Deinen Willen die Expedition erfolgreich zu beenden. | *Imagine you are an adventurer on your way to the north pole. You are writing a chapter of your travel diary, in which you emphasize your willingness to successfully bring the expedition to an end, despite low supplies and the bitter cold.* |
| Stell Dir vor, Dein Partner findet in Deinem Handy einen Nachrichtenaustausch mit einer/m potentiellen Geliebten. Ihr geratet in einen Streit und Du versuchst Dich zu rechtfertigen. | *Imagine your partner finds text messages with a potential lover on your mobile phone. You get into an argument and you are trying to explain yourself.* |
| Stell Dir vor Du bist Bundestagsabgeordneter der Grünen. Du hältst eine Rede über die Herausforderungen, die durch den Klimawandel entstehen, und die richtigen Maßnahmen, um eine katastrophale Erderwärmung aufzuhalten. | *Imagine you are a member of the parliament from the Green party. You are giving a speech about the challenges posed by climate change and the right measures to stop a catastrophic global warming.* |
| Stell Dir vor Du bist ein Strafverteidiger und vertrittst vor Gericht einen Bankräuber. Du hältst ein Plädoyer für den Angeklagten, der | *Imagine you are a defense attorney, and you are representing a bank robber at court. You are making a summation for the* |

| | |
|---|---|
| den Überfall begangen hat, um seiner Frau eine lebensrettende Operation zu finanzieren. | *accused, who committed the robbery to be able to pay for a life-saving surgery for his wife.* |
| Stell Dir vor Du bist Teil eines Höhlenforscherteams. Ihr werdet als erste Menschen eine neu entdeckte, steinzeitliche Höhle betreten. Vor der Expedition erklärst Du einem Fernsehteam, was die potenzielle Entdeckung neuer Höhlenmalereien für die Frühgeschichte der Menschheit bedeuten kann. | *Imagine you are part of a group of cave explorers. You will be the first humans to explore a newly discovered stone-age cave. Prior to the expedition, you speak to a tv team and explain to them what the potential discovery of new cave paintings would mean for understanding early human history.* |
| Stell Dir vor Du bist ein unbekannter Läufer, der total überraschend den zweiten Platz im Berlin-Marathon belegt. Die Medien bezichtigen Dich daraufhin des Dopings. Du versuchst die Öffentlichkeit in einem Pressestatement von Deiner Unschuld zu überzeugen. | *Imagine you are an unrenowned runner who totally surprisingly finishes the Berlin marathon in second place. Afterwards, the media accuse you of doping. In a press statement, you try to convince the public that you are innocent.* |
| Stell Dir vor Du bist ein deutscher Diplomat auf dem Weg zu einem wichtigen EU-Gipfel. Dort wirst Du gebeten darzustellen welche politischen Maßnahmen Du zur Eindämmung der „Flüchtlingskrise“ als besonders sinnvoll ansiehst. | *Imagine you are a German diplomat on your way to an important EU summit. There, you will be asked to outline political measures you consider suitable for containing the “refugee crisis”.* |
| Stell Dir vor Du bist ein sozialdemokratischer Politiker im Deutschland des 19. Jahrhunderts. Du bekommst die Chance vor dem Kaiser zu sprechen und ihn zu überzeugen die Rechte des Parlaments auszubauen um ein moderneres und gerechteres politisches System zu formen. | *Imagine you are a social democrat politician in 19th century Germany. You get the chance to speak to the Kaiser and convince him to strengthen the rights of the parliament to create a more modern and fairer political system.* |
| Stell Dir vor Du bist ein Bergarbeiter. Während eines Unglücks wird Du mit anderen Kumpels in einem Stollen eingeschlossen. Du denkst über die Gründe | *Imagine you are a miner. During an accident, you and some co-workers get trapped in a tunnel. You are reflecting on* |

| | |
|---|---|
| des Unfalls nach und taxierst eure Chancen gerettet zu werden. | *what had caused the accident and try to estimate the chances of getting rescued.* |
